# Shifts in nutrient allocation in a gift-giving butterfly: A hidden consequence of water balance?

**DOI:** 10.1101/2025.07.17.665443

**Authors:** Chloé Chabaud, Natasha Tigreros

## Abstract

As climate change intensifies drought, understanding how animals maintain fitness under water stress is key to predicting species persistence. Animals use diverse behavioural and physiological adjustments to avoid dehydration. However, the physiological and fitness costs of these mechanisms are often overlooked, despite their potential to shift nutrient acquisition and allocation. We hypothesized that maintaining water balance, through increased water intake and/or decreased water loss, leads to nutrient shifts and trade-offs in *Pieris rapae* butterflies. In this species, females receive a protein- and water-rich nuptial gift (NG), known to enhance fecundity and possibly mitigate dehydration. We quantified the impact of dry conditions on female hydration and fitness, using stable isotopes to trace nutrient allocation to storage, fecundity, and catabolism. We found that the NG, combined with reduced respiratory water loss, contributed to maintaining female water balance in dry conditions. Importantly, while dry environments did not impact potential fecundity, nutritional shifts and trade-offs that could affect long-term fitness were evident: females allocated more lipids to eggs at the expense of long-term storage, while reducing catabolism of NG-derived leucine. This interplay among water balance, nutrient allocation, and fitness emphasizes the importance of linking water balance mechanisms with broader nutrient-use strategies under environmental stress.

## 1 INTRODUCTION

Water is an essential resource for all living organisms, comprising more than 60% of body mass in most terrestrial animals. Maintaining adequate water balance is critical to animal fitness, supporting key physiological functions such as thermoregulation, digestion, and cellular homeostasis. However, availability of free-standing water is often limited and increasingly variable as extreme weather events—such as droughts—intensify in some regions due to climate change [1,2]. To avoid dehydration, animals can adjust their behaviour and physiology to increase water intake (e.g. through consumption of preformed water and food) and reduce water losses via cutaneous evaporation, excretion, or respiration (e.g. [3,4]). These water balance mechanisms have been extensively studied, particularly in arid-adapted species or in the context of climate change [5]. Yet relatively little attention has been given to their potential costs or constraints, even though such mechanisms are often linked to processes of acquisition and metabolism of other nutrients. Understanding how organisms balance the benefits and costs of water balance mechanisms is critical for predicting whole-organism responses and resilience to changing environments [6].

When animals obtain water through their diet, water balance becomes a key factor influencing their dietary preferences and foraging activity. Animals can regulate water intake by selectively feeding on foods with different preformed water and energy content [7,8]. For example, sap-sucking insects choose between feeding on nutrient-rich phloem or water-rich xylem, depending on their water versus energy requirements [9]. When choosing among diets is not an option, animals can increase food intake until their water needs are met or vice versa [10]. A consequence of such a strategy, however, is that it can result in excessive intake of other food components and thus affect the optimal expression of other functions [11]. While studies have typically focused on how organisms prioritize sources of caloric intake (e.g., protein vs. carbohydrates), limitations in water availability may similarly lead to nutritional reallocation and trade-offs critical to overall fitness.

Nutritional shifts may also arise when the mechanism to maintain water balance involves changes in metabolic processes, such as metabolic water production. The production of metabolic water through the oxidation of dietary macromolecules is a widespread strategy for terrestrial animals to avoid dehydration [12–15]. For example, birds and lizards under water stress are known to increase proteins oxidation (e.g. [16–18]), as protein catabolism produces more metabolic water than the oxidation of other macromolecules relative to the energy obtained, in part because it also releases bound water [19]. Other invertebrate species rely more heavily on lipid or glycogen catabolism (e.g. [20–22]). However, diverting lipids or protein for water production may reduce availability for other vital functions, such as reproduction, leading to physiological trade-offs. Thus, while their oxidation can provide a vital water source during dehydration, it comes at the cost of potentially impairing other aspects of fitness.

In addition to acquiring water metabolically, animals can conserve water by reducing their metabolic rate. In terrestrial environments, gas exchange through the respiratory system is a major route of water loss, as respiratory surfaces must remain moist to allow efficient diffusion of gases. Higher metabolic demands increase the rate and volume of gas exchange, thereby elevating evaporative water loss [23]. Consequently, lowering metabolic rate can be an effective water-saving strategy. This adaptation has been documented in desert birds along aridity gradients [24], as well as in mammals [25] and insects, which limit water loss via discontinuous gas exchange [26]. While reducing metabolic rate helps conserve water, it can result in significant fitness costs. For instance, reproductive success may decline due to slower growth rates and reduced fecundity [27]. Similarly, immune function can be compromised because of the reduced energetic investment associated with a lower metabolic rate [28].

Maintaining water balance is particularly challenging for insects. Due to their small body size and high surface-to-volume ratio, insects are predisposed to rapid water loss and desiccation [5,29,30]. Despite these constraints, insects have evolved remarkable adaptations to prevent dehydration. In some species, mating itself may offer a unique opportunity to supplement water intake through nuptial gifts—nutrient donations that females receive during copulation. These gifts, often rich in both water and nutrients [31], may serve a dual function in water-limited environments: providing paternal investment and mitigating female dehydration. Previous studies have shown that nuptial gifts are particularly rich in proteins, providing females with essential amino acids for egg production [32]. Marshall (1982) first hypothesized that butterfly nuptial gifts could provide both nutrients and water, particularly in arid environments [33]. Subsequent research has supported their potential hydration benefits across insects. For example, water-deprived *Gryllodes sigillatus* crickets maintained offspring production comparable to well-hydrated counterparts [34], and female moths that mated with water-supplied males showed increased longevity, suggesting water transfer during copulation [35]. Behavioural studies further reinforce this, with water-deprived female *Callosobruchus maculatus* beetles mating more frequently, suggesting that opportunities for hydration influence reproductive decisions [36,37]. Thus, nuptial gifts provide a great opportunity to examine the links between water balance, nutrient use, and the fitness costs of facing environmental stress.

In this study, we test the hypothesis that nuptial gifts in *Pieris rapae* butterflies help females cope with water stress while shifting the allocation of nutrients to other functions. *P. rapae* males transfer a nuptial gift (also known as a spermatophore) during mating. This gift is rich in sugars, nitrogen and water [38–40] and is known to substantially enhance female fecundity and longevity [41]. *P. rapae’*s wide geographic distribution [42] exposes it to diverse humidity levels and frequently fluctuating water availability, making their potential for utilizing nuptial gifts for both water and nutrients particularly relevant. To understand how nuptial gifts contribute to maintaining water balance, we tested the impact of dry conditions on hydration and fitness of mated versus virgin females. We then examined nutrient shifts and trade-offs using ^13^C-stable isotope to trace the allocation of lipids and a nuptial gift-derived essential amino acid to fecundity, catabolism and storage [43]. By integrating fitness and physiological measurements with real-time stable isotope analysis, our study provides a powerful approach to understand how organisms deal with water-limited environments.

## 2 MATERIAL & METHODS

### 2.1 General rearing and ^13^C-labeled nuptial gifts

*Pieris rapae* butterflies used for this experiment were from a laboratory colony that originated from a field population in Utah, USA. Butterflies were reared from egg to 5 days old (early 2^nd^ instar) on their host plant, *Brassica oleracea*. At this stage, larvae were transferred in pairs into 1 oz. plastic cups and fed *ad libitum* with a semi-synthetic diet [44,45]. This diet minimized variation in the larval nutritional environment (e.g., maintaining 4,2% nitrogen by dry mass) and ensured optimal conditions for the development of males so they produce a protein-rich nuptial gift. Throughout their development, butterflies were kept in a walk-in environmental chamber with LD 16:8 h photoperiod and 22°C at 50% relative humidity.

To examine how females use nutrients from the male nuptial gift, we isotopically enriched the male larval diet. At 10 days old, when males can be reliably identified based on visible testes, individuals were switched to either a ^13^C-leucine labelled or a “control” (unlabelled) diet until pupation [46]. ^13^C-leucine diet was made with an addition of 0.17 g L-Leucine (1-^13^C, 99%, Cambridge Isotopes Laboratories, Inc.) per 0.5 L diet and control diet with the same amount of L-Leucine (non-labelled, 99%, Alfa Aesar). Manipulation of the male larval diet resulted in a clear isotopic distinction between control and labelled nuptial gifts transferred to females; the δ^13^C signature of labelled gifts was significantly higher than that of both the control gifts and the female diet (Figure S1), confirming effective ^13^C enrichment. Since essential amino acids cannot be synthesized *de novo* by adults, we hypothesized that they would be preferentially allocated to reproduction over metabolism, given their limited availability and importance for reproductive success.

To mate females with either ^13^C-labelled or control males, newly emerged females were transferred to a greenhouse (14:10 L:D cycle) and placed in a large cage (15.7”D x 15.7”W x 23.6”H) with a similar number of males from each diet. To standardize nuptial gift investment, we only included virgin males [47] and individuals that were four days old or younger. Mated females were brought back to the lab to experience different humidity levels (see below).

### 2.2 Humidity treatments

After mating, females were randomly assigned to one of two humidity treatment: Dry (35% RH) or Wet (65% RH). These treatments reflect natural humidity levels encountered across *P. rapae*’s range [42], with wet being ideal, non-desiccating conditions [48] whereas dry is a challenging, desiccating environments [49]. Females were kept for 48 hours in individual cages (9”L x 9”W x 11”H) within controlled greenhouses equipped with Inkbird humidity controllers and Dreyoo humidifiers.

To control for water-saving strategies independent of nuptial gift use, virgin females were included in both humidity treatments. To standardize energy intake and ensure a uniform need to mobilize nuptial gift resources regardless of humidity effects, all females were hand-fed 5µL of a 10% sucrose solution 24h after entering the humidity treatment.

### 2.3 Measuring the benefits of nuptial gifts on water balance and fitness

To assess the benefits of nuptial gifts for females experiencing dehydrating environments, we compared the water balance and survival of virgin and mated (with a nuptial gift) females when experiencing either a wet or dry environment. We tested whether female water balance—measured as haemolymph osmolality and whole-body water content—was influenced by environmental humidity, mating status (nuptial gift receipt), or their interaction. Measuring haemolymph osmolality helps determine the concentration of solutes in the insect’s circulatory fluid, which can increase during dehydration as water is lost from the body. However, osmolality can also be influenced by other factors, so it is important to consider those two complementary measurements together [50]. We predicted that dry conditions would increase osmolality and reduce water content, while nuptial gifts would buffer water loss by lowering osmolality and/or increasing water content. To collect haemolymph for osmolality measurements, females were first immobilized by placing them in a freezer (−20°C) for three minutes to facilitate handling. Then, a minimum 1 µL of haemolymph was extracted by piercing the thorax at the mesothorax-metathorax junction with a fine microcapillary tube. Samples were transferred to a microcentrifuge tube, diluted with 10 µL of stable saline solution (osmolality = 285.8 ± 1.7 mOsm/kg), and stored frozen until analysis. Osmolality was measured using a Wescor Vapro Model 5600 osmometer. If more than 1 µL was collected, the sample was duplicated to assess repeatability, and values were adjusted for dilution. Body water content (%) was calculated gravimetrically as: (Wet Mass-Dry Mass)/(Wet Mass)*100. Females were weighed before haemolymph collection to obtain the wet mass using a high precision scale (Sartorius Quintix65-1S), and again after being killed and dried at 50°C for 48h to obtain dry mass.

Survival of mated and virgin females under the two different humidity conditions was measured until females were sacrificed after 48 hours of humidity treatment for water balance measurements, and any mortality occurring before 48 hours was recorded to determine whether dry conditions differentially affected short-term survival between mated and virgin ones.

### 2.4 Evaluation of nuptial gift allocation under dry conditions and potential costs

To assess the potential costs of coping with dry conditions, we focused on both nuptial gift consumption and its allocation to reproduction and metabolism. We first quantified nuptial gift consumption over 48 h in both environments by comparing the dry mass of the remaining nuptial gift under dry versus wet conditions. While the initial mass of these nuptial gifts was not quantifiable, a pilot study indicated that nuptial gifts contain approximately 82% (3-4 mg) immediately after mating. We then assessed how nuptial gift-derived nutrients were used, specifically the allocation of leucine to either egg production or oxidation. To evaluate potential fecundity, we dissected mated females to remove their eggs and ovaries, dried the tissues at 50□°C for 48 h, and recorded the dry mass.

At 48h post-mating, we sampled breath isotopes in mated females using a CRDS (G212-I, Picarro, Santa Clara, CA) to quantify CO_2_ production linked to gift-derived ^13^C-leucine [51]. Male *P. rapae* were reared on an artificial diet enriched with ^13^C-leucine (as described above), resulting in nuptial gifts that were isotopically distinct from other female food sources (larval diet and beet sugar nectar, δ^13^C = -26.5‰, Figure S1). Each female was placed in a 10mL respirometry chamber for 10 min, after flushing it with air previously deprive of CO_2_ and water (using a column containing Soda Lime and drierite). Following the incubation, the chamber air was directed at 30 ml/min into the G212-i CRDS stable carbon isotope analyser, with data recorded at 0.5 Hz using PICARRO software. CO_2_ and H_2_O measurement provided estimates of metabolic rate (VCO_2_ in ml/min) and respiratory water loss. We chose to take measurements at 48h based on prior work in *P. rapae* and similar species that indicated females start digesting the NG within the first hour after mating [52], and that usage of nuptial gift-derived nutrients starts peaking on day two after mating in close-related species [53].

To compare how females under different RH% environments used nutrients (including nuptial gift-derived leucine), we dissected the abdomen of each female to separate eggs/ovaries from the fat body and residual nuptial gift. Tissue samples were dried at 50°C for 48 hours (VWR 1300U laboratory oven), homogenized, and weighed. These samples were then loaded into tin capsules for δ^13^C measurement using a Picarro G2121-i stable isotope analyser. We obtained ^13^C concentrations of both control and labelled tissues expressed in δ^13^CVPDB [54] and subsequently calculated the atom percent (AP) and proportional ^13^C allocation to metabolism, eggs, and somatic tissues [54]. Because environmental conditions are known to influence natural stable isotope signatures, we first examined variation among control females (mated to non-labelled males) exposed to dry or wet conditions. In addition to tracing labelled leucine, we used the δ^13^C signatures of control females to assess endogenous nutrient allocation patterns. Because lipids are ^13^C-depleted relative to proteins and carbohydrates [55], shifts in δ^13^C across tissues in control individuals provided insight into how water availability may influence macronutrient use. To account for this background variation in control females, we used Atom Percent Excess (APE), defined as APE = AP(label) – AP(control) [56], for subsequent analyses of leucine.

To analyse differences in relative allocation, AP values were converted into the mass of CO_2_ and ^13^CO_2_ in each pool (breath, eggs, fat body). For eggs and fat body, CO_2_ mass was calculated by multiplying the sample’s CO_2_ concentration by the total dry mass of the pool (normalized by the sample mass), and the ^13^CO_2_ mass was derived by multiplying this value by the AP of ^13^C (divided by 100). For breath, we integrated the metabolic rate over 24h to account for the timing of resource use from the nuptial gift, as females begin metabolizing it one day post-mating [53]. Finally, to assess individual differences in nutrient allocation, we calculated the total ^13^C allocated to each pool and determined the relative allocation by dividing each pool’s allocation by the total ^13^C allocation with the following calculation: 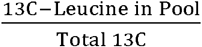. We hereafter refer to nuptial gift-derived leucine as the labelled leucine originating from the nuptial gift.

### 2.5 Statistical analysis

#### 2.5.1 For osmolality and body masses

We used linear mixed-effects models (lmer() from the lme4 package in R) to analyse osmolality and water content. The model included Humidity treatment, Female status (virgin or mated) and their interaction as fixed effects, with batch number as a random effect. Similarly, to compare egg production and the partial digestion of the nuptial gift in mated females, we modelled egg dry mass and residual nuptial gift dry mass as functions of Humidity treatment and total dry body mass (with batch as a random effect). Model assumptions (normality and homoscedasticity of residuals) were verified through visual inspection of residuals and Q-Q plots.

#### 2.5.2 For survival analysis

We first examined the survival rates of females under two different conditions: dry and wet. We performed a Mantel-Haenszel test [57] to evaluate the association between mating status (virgin vs. mated) and survival (survived vs. died) across the two different treatments (DRY and WET). The test was conducted using an exact method to account for the small number of butterflies that died during the 48h experimental period in some groups (fewer than 5 individuals in both mated and virgin groups under wet conditions).

#### 2.5.3 For isotopic measurements and metabolic rate

We first compared δ^13^C in eggs, fat body, and breath of control females to assess how relative humidity affected nutrient allocation in the absence of enrichment. We then compared isotopic signatures between females that received nuptial gifts from control (unlabelled) versus ^13^C-leucine–labelled males to test whether females allocated nuptial gift-derived leucine, an essential amino acid, to all three pools. We used linear mixed-effects models with atom percent (AP) as the response variable, and nuptial gift label, sample CO□, and pool dry mass (or total female dry mass for breath) as predictors, including batch as a random effect.

Second, to test the effect of humidity on nuptial gift allocation, we analysed only females that received labelled nuptial gift. We used atom percent excess (APE = AP(label) – AP(control) [56]; as the response variable, with humidity treatment, pool mass, and sample CO□ as predictors (again, using total female dry mass for breath).

Third, we assessed each individual’s relative allocation of leucine by modelling the proportion of ^13^C in each pool relative to the total ^13^C. Finally, to test for trade-offs between allocation pathways, we modelled ^13^C allocation to one pool as a function of allocation to another, including an interaction with humidity treatment. Significance of fixed effects was assessed using Type II or III ANOVA (Anova function, car package).

Similar models were used to compare metabolic rate and evaporative water loss between dry and wet environments, with humidity treatment and female dry mass as covariates. Descriptive statistics are reported as means ± SE.

## 3 RESULTS

### 3.1 Benefits of nuptial gifts for water balance

Assessment of both haemolymph osmolality and body water content indicated that mated females were less dehydrated than virgin females when exposed to dry conditions. First we found a significant main effect of humidity treatment on haemolymph osmolality, with females from the dry environment exhibiting higher osmolality compared to those from the wet environment (χ^2^ = 10.62, df = 1, p = 0.001; Osmolality in Dry = 460±8 mOsm.kg-1, Wet = 425±6 mOsm.kg-1; Figure 1A). Second, the effect of humidity treatment on female’s total body water content was dependent on the female mating status (χ^2^_humidity*mating_ = 4.65, df = 1, p = 0.03; Figure 1B); while virgin females showed reduced water content in the dry environment relative to the wet (mean = 59.7±1% in dry conditions; 62.8±0.8% in wet conditions), mated females maintained similar water levels across both environments (mean = 65.8±0.6% in dry; 65.7±0.7% in wet). Female mass, included as a covariate, also influenced water content (χ^2^_mass_ = 23.89, df = 1, p<0.001).

**Figure 1:**
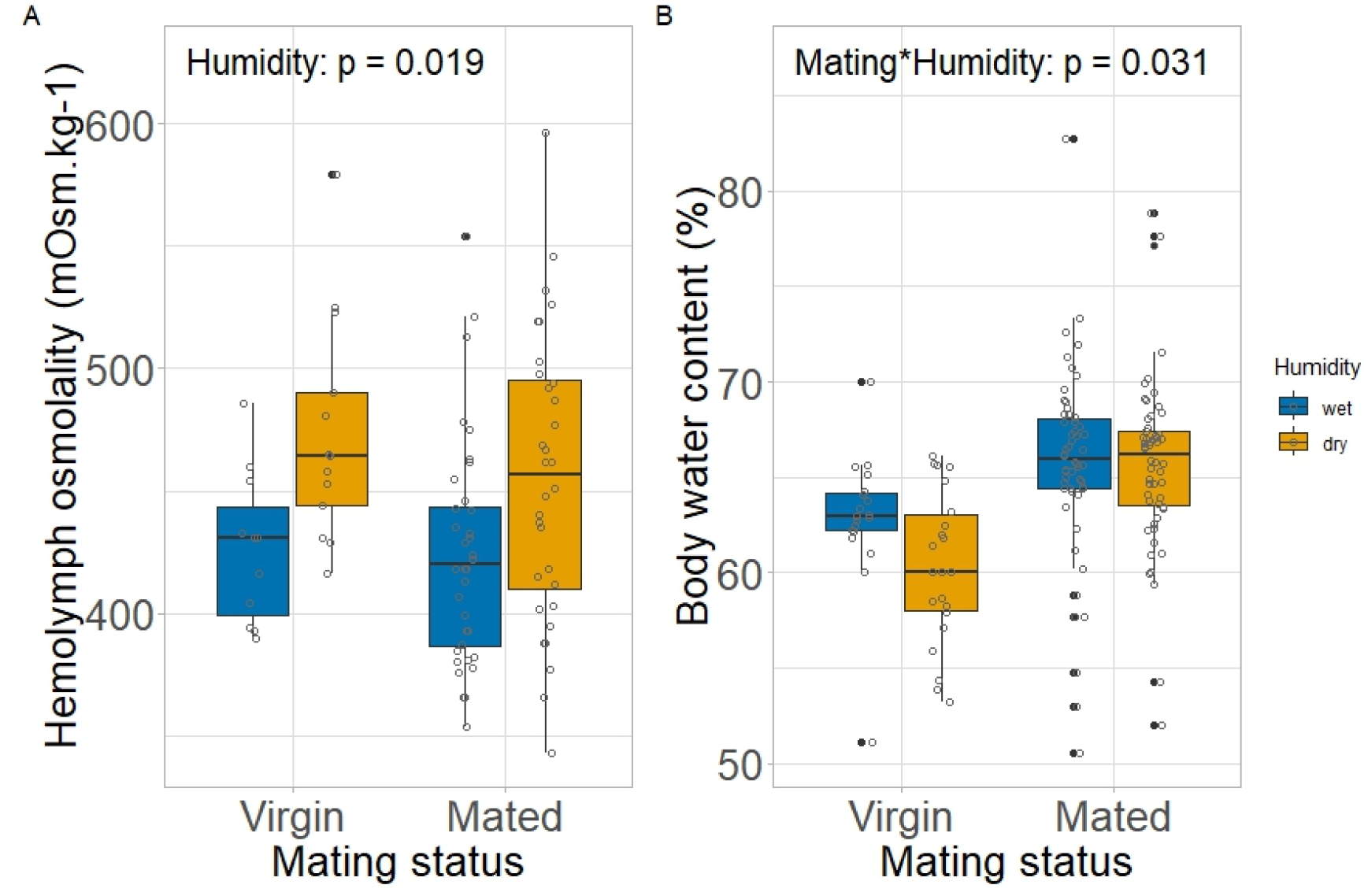
Effect of mating status (virgin vs. mated) on water balance in *P. rapae* females exposed to either a wet (blue, 65% RH) or dry (orange, 35% RH) environment. Water balance was estimated using two complementary variables: (A) Haemolymph osmolality (mOsm·kg□^1^) and (B) body water content (%) (B). Higher osmolality indicates dehydration, particularly when coupled with water loss. Boxplots show the median, interquartile range, and data spread; individual data points are shown as circles.

### 3.2 Effect on fitness

Survival analysis results indicated that mated females had significantly higher odds of survival than virgin females across humidity environments (Mantel-Haenszel test, p-value = 0.05, Figure S3). Although the survival benefit of mating tended to be greater in dry conditions (a 10.7% increase in survival in dry versus 9.9% in wet conditions), this difference was not statistically significant (p = 0.09). We also found no significant difference in female potential fecundity—measured as total dry mass of oocytes 48 hours after mating—between mated females from dry and wet environments (p = 0.46, Figure S3).

### 3.3. Water balance mechanisms

#### 3.3.1 Respiratory water loss

As expected, we found that respiratory water loss and metabolic rate were positively correlated, with higher VCO_2_ values associated with greater respiratory water loss (p = 0.006, Figure S4). Importantly, females in the dry environment, compared to those in the wet environment, exhibited lower metabolic rate (χ^2^ = 4.12, df = 1, p = 0.04; Figure 4A) and significantly reduced water loss through breath (χ^2^ = 9.31, df = 1, p = 0.002, Figure 4B), with female dry mass included as a significant covariate for metabolic rate (χ^2^ = 16.12, df = 1, p<0.001).

#### 3.3.2 Nuptial gift consumption

We found no support for our hypothesis that females in dehydrating environments might accelerate consumption of nuptial gifts to obtain more water. Comparison of nuptial gift remains–their dry mass 48h after mating–indicated that there was not a significant difference in consumption by females under dry versus wet environments (p = 0.89, Figure S2).

### 3.4 Nutrient utilization and trade-offs

#### 3.4.1 Female use of lipids

Comparison of δ^13^C in control females (mated to unlabelled males) indicated that females in the dry treatment had more negative δ^13^C in eggs (p = 0.03, Figure S5) and less negative δ^13^C in the fat body (p = 0.05, Figure S5) compared to females in wet treatment. Given that lipids are naturally ^13^C-depleted compared to proteins and carbohydrates [55,58], this suggested that females in dehydrating environments incorporated more lipids into eggs but less into storage. Comparison of δ^13^C_breath_ among treatments revealed no evidence of altered lipid catabolism under dehydrating conditions (p = 0.76).

#### 3.4.2 Female use and allocation of NG-derived leucine

Comparison of δ^13^C from females receiving a control versus a ^13^C-leucine labelled nuptial gift revealed that females deposited this essential amino acid both into eggs (χ^2^ = 262, df = 1, p<0.001, Figure 2A) and into storage (χ^2^ = 23.2, df = 1, p<0.001, Figure 2B). Also, breath analysis further revealed significant differences in δ^13^C of both female groups (χ^2^ = 217, df = 1, p<0.001, Figure 2C), indicating that females used NG-derived leucine as a metabolic fuel.

**Figure 2:**
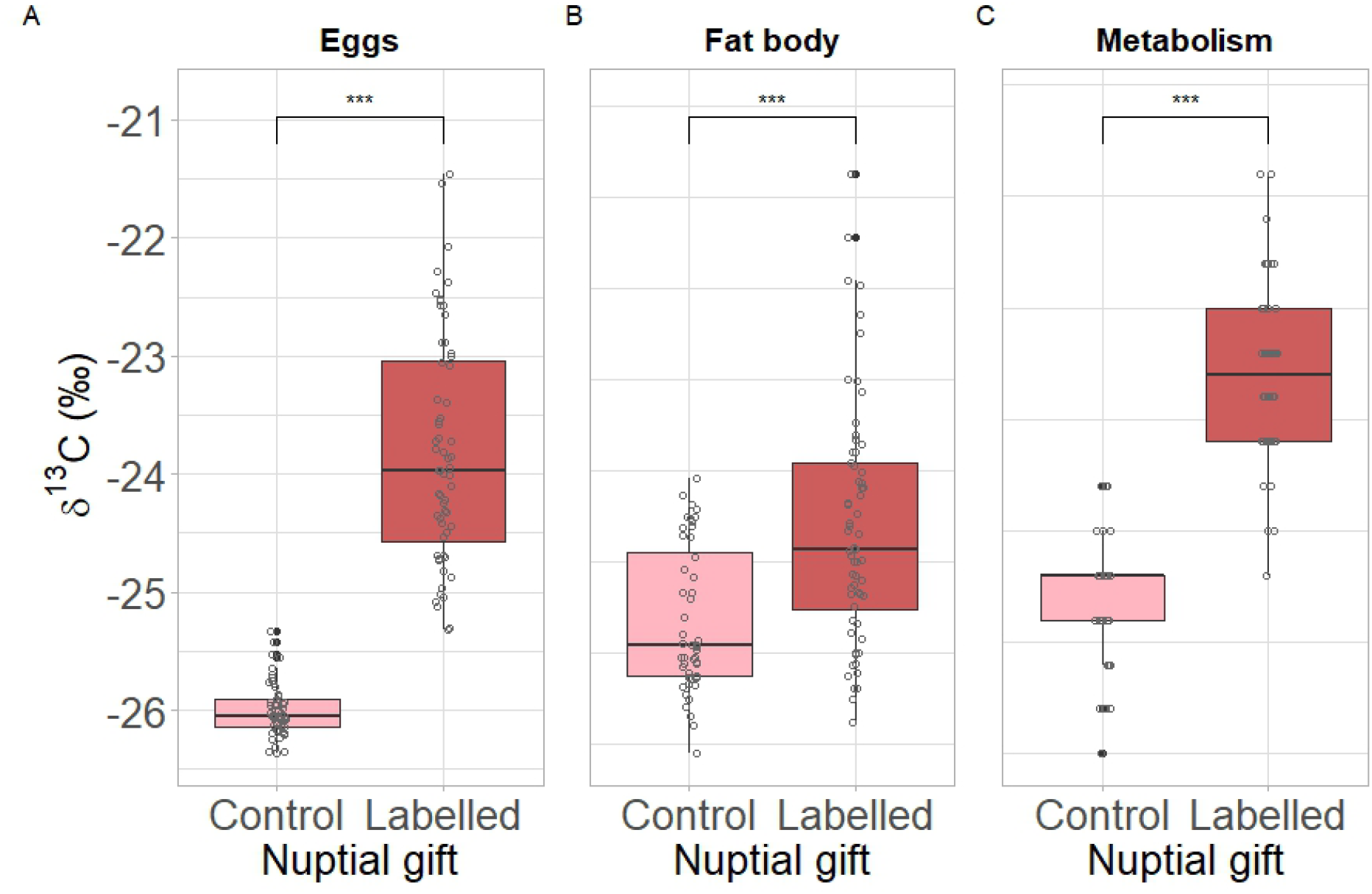
The δ^13^C values (‰) in different body parts of *P. rapae* females that received either a control nuptial gift (pink), or a labelled nuptial gift from a male raised on ^13^C-labelled leucine diet (red). Values are shown in eggs (A), fat body (B), and breath (C) to demonstrate allocation of nuptial gift-derived leucine to reproduction, storage and metabolism. Boxplots show the median, interquartile range, and data spread; individual data points are shown as circles. Less negative δ^13^C values indicate higher tracer content. ***p < 0.005.

Relative humidity did not significantly affect the absolute amount of NG–derived leucine allocated to eggs (p = 0.25; Figure 3A) or stored in the fat body (p = 0.45; Figure 3B), but did lead to a marginally significant effect on leucine catabolism with females in the dry treatment increasing leucine oxidation compared to those in the wet treatment (breath APE: Estimate = 0.0011 ± 0.0005, p = 0.054; Figure 3C). At the same time, when comparing the relative allocation of total recovered NG-derived leucine, females in dry environments allocated a smaller share for catabolism (calculated as the ratio of ^13^C_breath * 24h_ / ^13^C_total recovered_) than those in wet environments (χ^2^ = 5.37, df = 1, p = 0.02; Figure 3D); this test accounted for the significant effects of female dry mass (χ^2^ = 22.98, df = 1, p < 0.001). Finally, no significant differences were observed in the relative allocation of leucine to eggs (p = 0.51) or to the fat body (p = 0.77) between humidity treatments.

**Figure 3:**
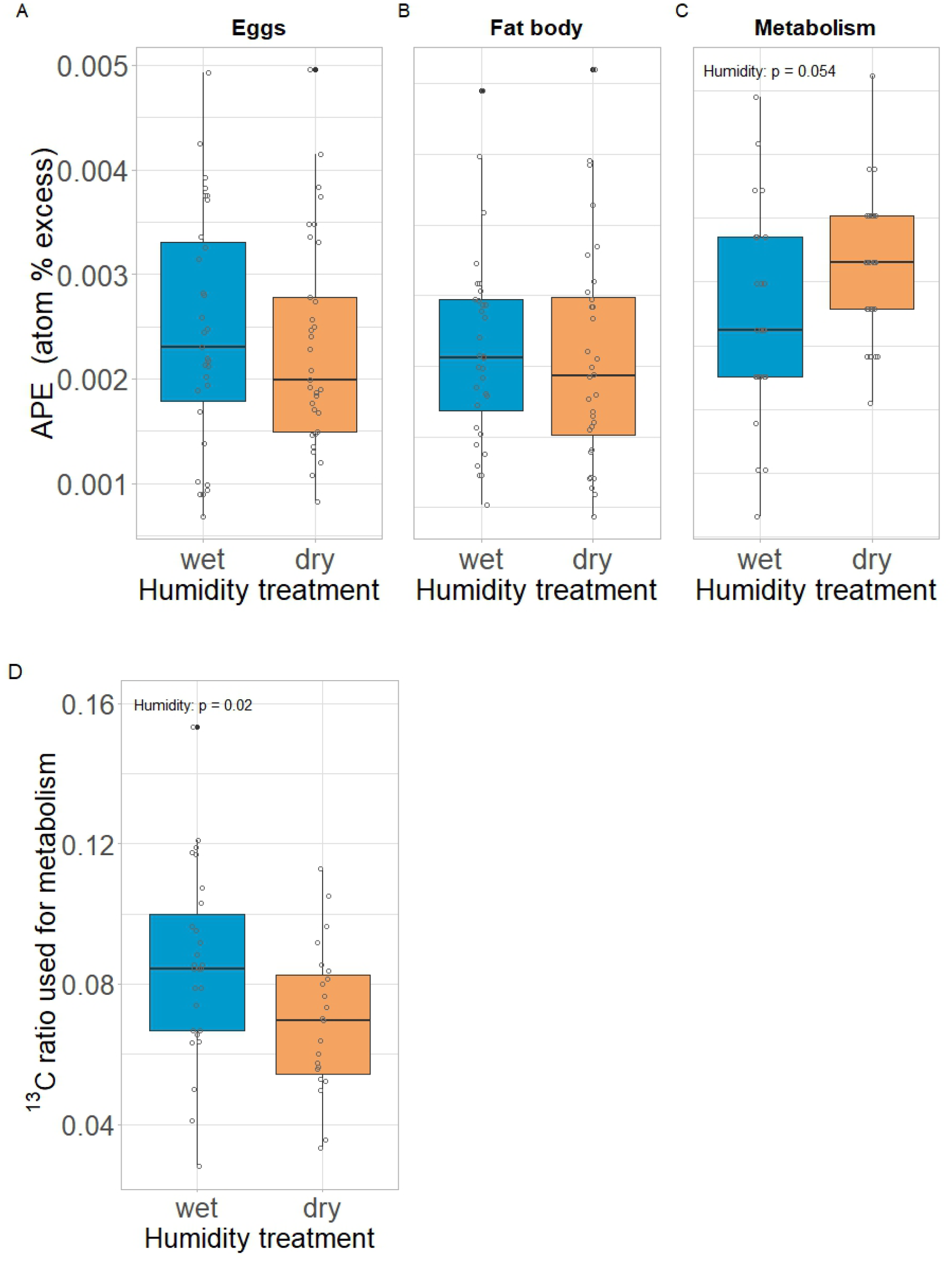
Difference in atom percent excess (APE, enrichment in ^13^C) in *P. rapae* females that received a labelled nuptial gift from a male raised on ^13^C -leucine diet and experienced either a wet (blue, 65% RH) or a dry (orange, 35% RH) environment after mating. Values are shown in eggs (A), fat body (B) and breath (C) to show different allocation of nuptial gift-derived leucine to reproduction, storage and metabolism under two humidity conditions. The ratio of ^13^C-leucine used as fuel for metabolism in 24h over the total female usage of ^13^C-leucine is shown in D. Boxplots show the median, interquartile range, and data spread; individual data points are shown as circles.

**Figure 4:**
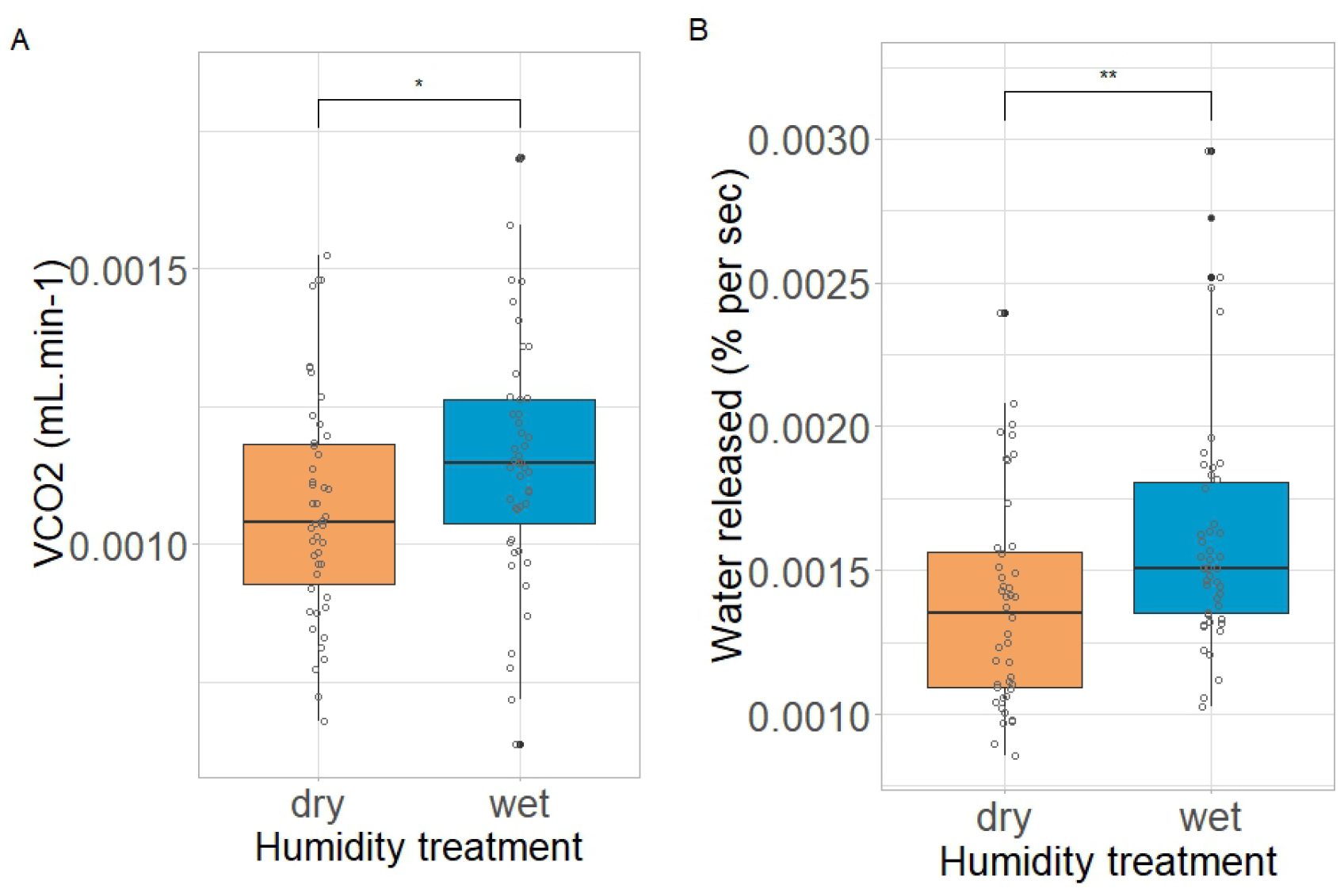
Metabolic rate (VCO_2_, A), and water loss in breath (% per second, B) in *P. rapae* females that experienced either a wet (blue, 65% RH) or a dry (orange, 35% RH) environment in the 48h following mating. Boxplots show the median, interquartile range, and data spread; individual data points are shown as circles. Statistical significance between the two humidity groups was assessed using the Wilcoxon rank-sum test; *p < 0.05; **p < 0.01.

Although we found evidence that the allocation of NG–derived leucine to metabolism could directly reduce its availability for other functions, this was independent of humidity treatment. Overall, females that oxidized a greater proportion of leucine also allocated more to egg production (χ^2^ = 4.19, df = 1, p = 0.04), with no significant effect of humidity (Estimate = 0.97). In contrast, leucine oxidation was negatively associated with allocation to the fat body (χ^2^ = 17.34, df = 1, p < 0.001), again independent of humidity treatment (Estimate = –1.97).

## 4 DISCUSSION

Maintaining effective water balance is crucial for organisms in water-limited environments. However, the costs and trade-offs that might arise when increasing water intake and/or reducing water loses remain largely underexplored, limiting our ability to fully understand and predict how organisms respond to increased drought and temperature extremes [5,6]. In this study, we addressed this issue by investigating how water balance, enhanced by the nuptial gift received during mating, affects female nutrient use and fecundity in *Pieris rapae* butterflies experiencing dehydrating environments.

### 4.1 Impact of the nuptial gift on females’ hydric state and fitness

Although primarily seen as a form of paternal investment [32], nuptial gifts can also play a crucial role in enhancing stress tolerance by supporting female water balance, especially in species experiencing variable conditions [59]. Yet, evidence for the potential hydration benefits of nuptial gifts has thus far been primarily indirect, relying on studies examining female mating frequency in relation to water availability or comparing offspring production under hydrated versus dehydrated conditions [34,36,37]. Our study provides the first direct physiological evidence that male nuptial gifts help females maintain water balance, demonstrating that a single gift increased female body water content by over 5% under dry conditions.

However, body water content alone does not fully capture hydration status. To better assess the physiological effects of mating, we also measured haemolymph osmolality, which can rise not only from water loss but also from the absorption of solutes such as amino acid or salts during nutrient mobilization [13]. While osmolality increased under dry conditions in both mated and virgin females, only virgin ones exhibited a concurrent decrease in body water content—indicating true dehydration. In contrast, mated females retained more water despite the osmolality increase, suggesting that elevated solute levels—possibly from nutrient uptake via the nuptial gift or active osmoregulatory processes—rather than water loss, contributed to the elevated osmolality [50]. This supports the idea that the nuptial gift buffered females against dehydration and helped them maintain cellular hydration under dry conditions.

Unexpectedly, nuptial gifts offered no measurable benefit for female short-term survival or fecundity in dehydrating environments. Although our experimental treatments were impactful—dry conditions increased mortality and mated females showed better overall survival due to the nuptial gift’s nutritional benefits [31] — the physiological hydric benefits of the gifts did not translate into a clear fitness advantage. This may stem from the short-term nature of our fitness estimates, as we measured potential fecundity and early survival (within 48 hours of mating) instead of realized fecundity and total lifespan. Fitness effects may become evident later, during egg maturation or through differences in egg water content and subsequent hatching success. Altogether, these results highlight the value of physiological measures, which can reveal early and subtle changes in female condition before they become evident in fitness outcomes.

### 4.2 Water balance mechanisms and nuptial gift use

We found that a single *P. rapae* nuptial gift, containing over 80% water by wet mass, could provide approximately 3–4□mg of preformed water. This amount closely matches the body water gain observed in mated females under dry conditions, suggesting that the direct intake of the nuptial gift’s water could account for a significant portion of the observed hydration benefit. However, the mechanism by which *P. rapae* utilize the gift’ water appeared to be complex, as females did not alter the consumption rate of it in dry conditions. This suggests that females do not fully control the rate of water (and nutrient) uptake; instead, this process may be largely influenced by the male ejaculate composition [60].

The observed hydration gain, despite the lack of increased water intake via consumption, suggests that complementary water balance mechanisms, including water loss reduction and metabolic water production, may explain how females benefit from the gift. Indeed, females in dry conditions significantly decreased metabolic rate, which thus reduced their respiratory water losses by almost 15%. This is a common physiological adaptation observed across diverse insect taxa for coping with water scarcity [23,61,62]. Interestingly, we also found some indication that the catabolism of nutrients derived from the nuptial gift could contribute to water balance in *P. rapae* females. Specifically, we observed a trend toward increased catabolism of nuptial gift–derived leucine under dry conditions. Protein (or amino acids) catabolism is an important mechanism for water generation in some vertebrate species [16– 18]. Although uricotelic insects excrete nitrogen with minimal water loss—suggesting amino acid catabolism could efficiently generate internal water—evidence for this strategy is notably absent within insect taxa. Instead, studies have shown that insects employ alternative strategies, favouring lipid metabolism [21] or relying on carbohydrate catabolism— particularly in desert-adapted species [20,62].

Interestingly, although there was a trend for a greater proportion of expired carbon in dry females to be derived from NG-derived leucine, a smaller share of their total recovered NG-derived leucine was allocated towards catabolism compared to those in wet environments. This shift was at least partly driven by the observed reduction in metabolic rate among females in dry conditions. Indeed, organisms across diverse taxa often reduce their metabolic overhead during periods of environmental stress. We see this in insects entering diapause [63], birds and mammals undergoing torpor [64], and even desert-dwelling amphibians decreasing metabolic activity to conserve both water and energy during dormancy [65]. Our results, therefore, indicate an integration of water-saving mechanisms with broader nutrient conservation strategies when facing challenging environmental conditions.

### 4.3 Shifts in nutrient allocation and trade-offs

Our study revealed strategic reallocation and trade-offs in the utilization of both lipids and NG-derived leucine by females in dehydrating environments. Although we found no evidence of increased lipid catabolism, females in dry environments increased lipid deposition into eggs while simultaneously reducing lipid storage in the fat body. This pattern is consistent with reproductive trade-offs commonly observed in insects, where limited resources are prioritized for immediate reproductive output at the expense of future reproduction [66,67]. However, increased lipid content does not necessarily improve egg quality, as some studies have shown that eggs richer in lipids have lower protein content, which can negatively affect hatching success [68]. Because lipids may originate from both the larval diet and the nuptial gift—and did not appear to contribute to metabolic water production—the observed lipid shifts likely reflect broader changes in how females manage internal resources under environmental stress. One possibility is that by allocating more lipids to eggs, females conserve protein and glycogen for their own somatic use, as these substrates generate more metabolic water during catabolism [19]—a potentially advantageous strategy in dry environments.

Females broadly allocated leucine across eggs, storage, and metabolic fuel. Importantly, they exhibited a clear allocation trade-off, prioritizing the nuptial gift-derived essential amino acid for immediate metabolic needs and reproduction over long-term storage. For essential amino acids, which cannot be synthesized, their immediate use for vital processes not only provides direct benefits but avoids costs associated with storing nutrients: energy for synthesis, maintenance, and potentially breakdown later [69]. Intriguingly, despite our prediction that water stress would alter nuptial gift resource allocation, leucine allocation trade-offs remained consistent across dry and wet environments. Our results therefore indicate that, for this essential amino acid, the female’s strategic prioritization of immediate reproduction and metabolic function overrides environmental water availability.

In summary, our findings indicate that nuptial gifts aid females in coping with dehydration by directly supplying water and by enabling flexible nutrient allocation decisions that optimize fitness under stress. Thus, the primary value of nuptial gifts in dehydrating environments may not be that of enhancing short-term fitness (survival or fecundity) but supporting the physiological adjustments necessary for reproduction to continue during stressful conditions. This crucial flexibility—evident in altered lipid allocation, selective nutrient catabolism, and changes in metabolic rate—likely underpins the ecological success of *P. rapae* in environments with variable water availability and illustrates a powerful intersection between mating interactions and stress physiology in shaping fitness outcomes under climate change.

### 4.4 Implications and future perspectives

Our results indicate that while nuptial gifts provide both water and nutrients, their use is shaped by physiological trade-offs that may influence future reproductive output or survival. Future research should investigate how these benefits manifest across the full reproductive lifespan of females, especially in terms of realized fecundity and offspring success during prolonged drought. It will also be important to explore the long-term consequences of altered nutrient allocation strategies—such as increased lipid investment in eggs—on both offspring quality and maternal survival in environments characterized by fluctuating water availability.

These findings also raise new questions about the adaptive value of nuptial gifts in water-limited environments. If gifts contribute to female hydration, their production may impose additional costs on males under hydric stress. This challenges the idea that water in nuptial gifts serves primarily as a manipulative tool to delay female remating [70,71], and instead suggests a potentially mutualistic role. Large gifts may be favoured for their dual function: extending the refractory period and providing water and other nutrients [32]. Future work could examine whether males adjust gift composition (e.g., water or protein content) in response to water availability.

Addressing these questions will help clarify how water balance, nutrition, and reproductive strategies interact under environmental stress. These mechanisms could influence species resilience under climate change, affecting not just individuals but also broader community dynamics where water availability is a limiting factor. Such insights are critical for predicting how organisms—and the systems they inhabit—will respond to increasing drought and climatic variability.

## Acknowledgements and funding

This work was supported by National Science Foundation (NSF-USA) grant IOS-2122282 to NT. We would like to thanks Madeleine Cahill and Gabriel Pearson for their help with colony maintenance and data collection as well as Anna Weinberg, Marnesha Jones and Goggy Davidowitz for insightful discussions and feedback.

