## Supplementary material for "Shifts in nutrient allocation in a gift-giving butterfly: A hidden consequence of water balance?"

**Figure S1: δ^13^C signature of control and labelled nuptial gifts, compared to the female larvae diet**

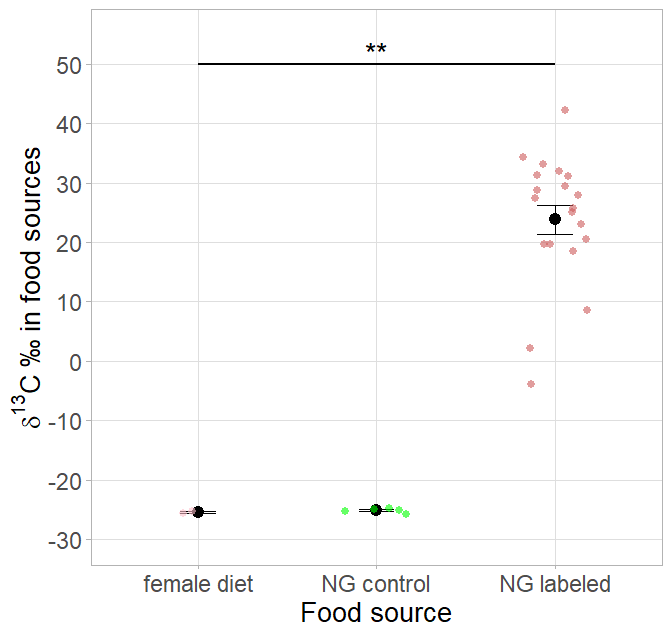

**Figure S1.** Plot showing the δ¹³C values across food sources available to *P. rapae* females. Nuptial gifts obtained from labelled males had significantly higher δ¹³C signatures than both the female larval diet and the nuptial gifts obtained from control males. **p < 0.01

**Figure S2: Nuptial gift consumption in dry and wet environments**

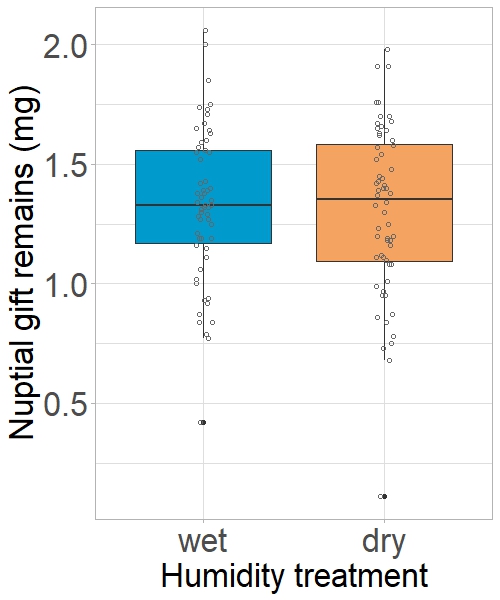

**Figure S2.** Boxplot showing the distribution of the dry masses of nuptial gift remains from *P. rapae* females that experienced either a wet (blue, 65% RH) or a dry (orange, 35% RH) environment for 48 hours after mating. This measure was used to assess nuptial gift consumption under different humidity conditions. Boxplots show the median, interquartile range, and data spread; individual data points are shown as crosses. No significant difference was observed between treatments.

**Figure S3: Fitness consequences of wet versus dry environments**

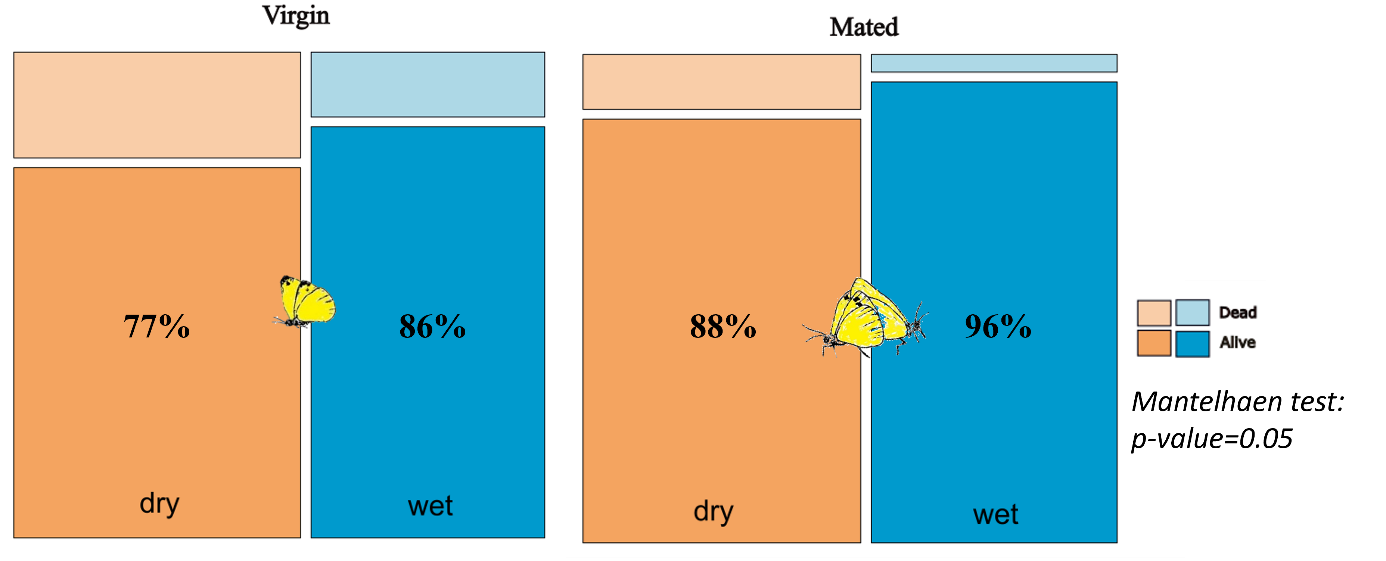

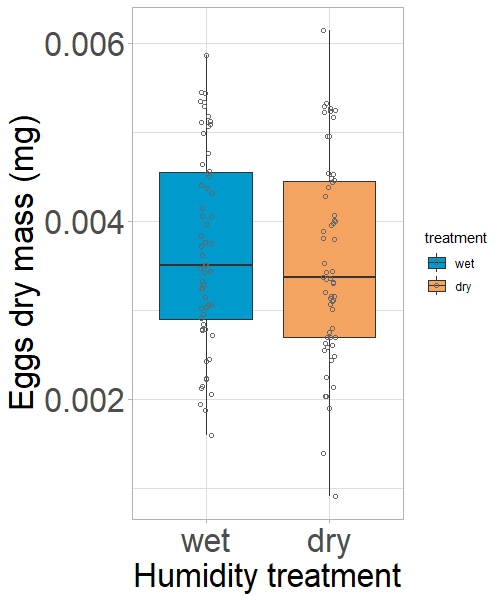

**Figure S3. (A)** Survival probability of *P. rapae* females during the 48-hour experimental period in either a wet (blue, 65% RH) or dry (orange, 35% RH) environment. The left panel shows virgin females; the right panel shows mated females. Survival was significantly lower in the dry environment overall (Mantel-Haenszel test, p = 0.05) **(B)** Potential fecundity of *P. rapae* mated females after 48 hours in either a wet or dry environment, measured as the dry mass of eggs and ovaries. No significant difference was observed between treatments.

**Figure S4**: **Relationship between metabolic rate and water loss**

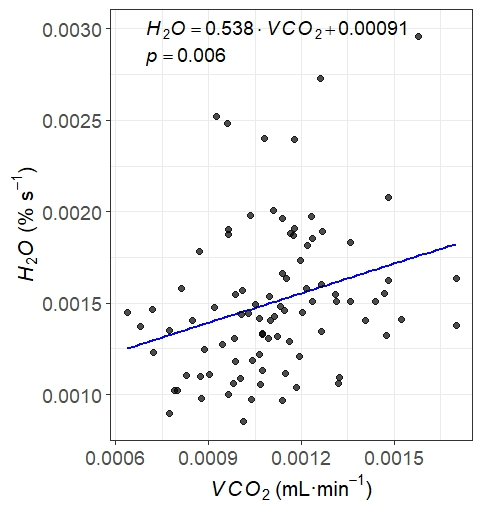

**Figure S4.** Relationship between V_CO2_ (mL/min) and water loss (% per second) during the 10-minute respirometry trial in *P. rapae* mated females. The solid line represents the linear fit. A Spearman correlation was used to assess the strength and direction of the relationship (ρ = 0.34, p = 0.00091).

**Figure S5:** **δ^13^C signature of eggs, fat body and breath from control females experiencing different environment**

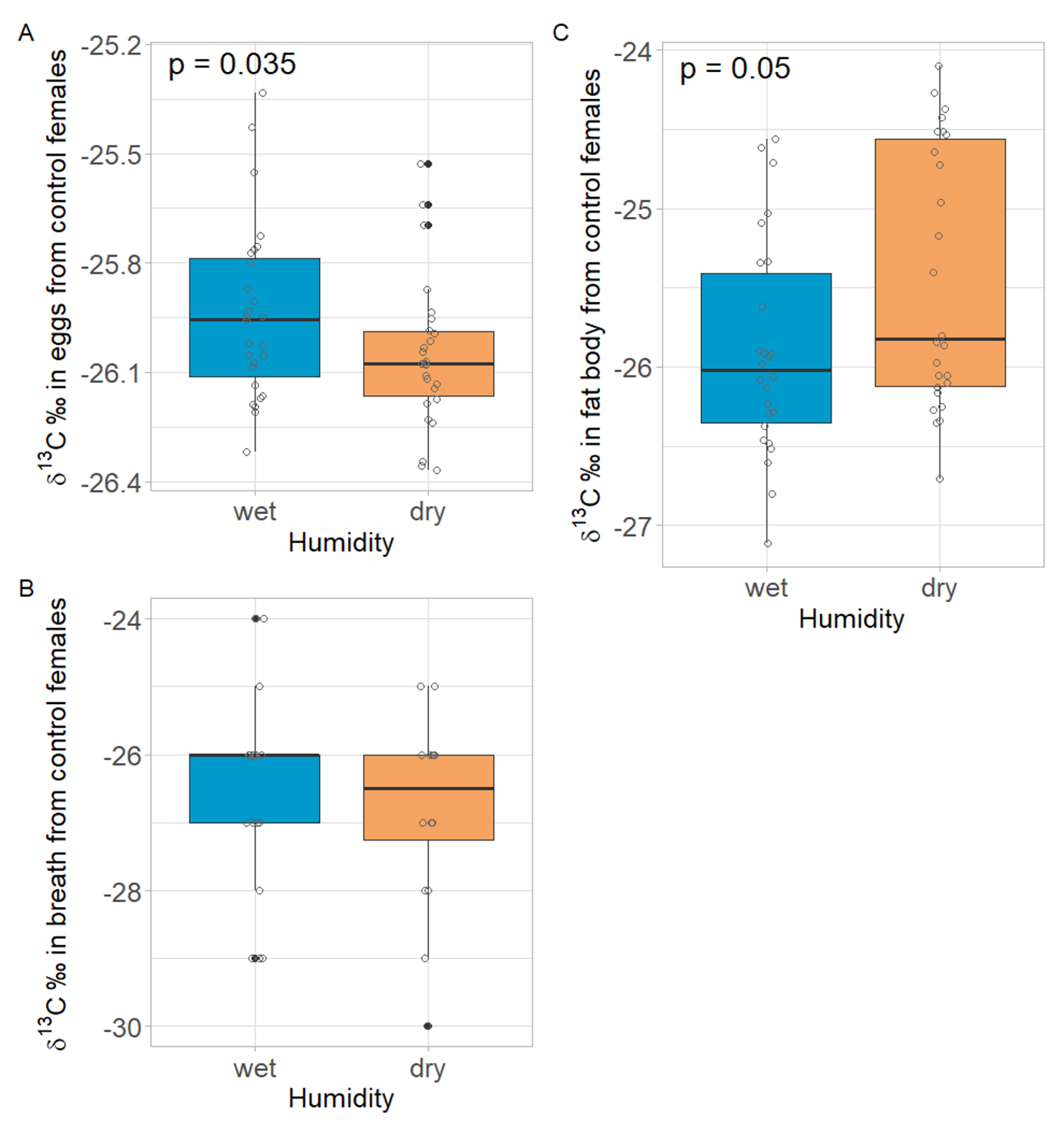

**Figure S5**: Isotopic signatures (δ¹³C) of control females mated with unlabelled males, reflecting the relative allocation of lipids versus proteins and carbohydrates across three physiological pools: (A) eggs (reproduction), (B) breath CO₂ (metabolism) and (C) fat body (storage). Because lipids are naturally ¹³C-depleted compared to proteins and carbohydrates, more negative δ¹³C values indicate greater lipid contribution. Under dehydrating conditions, females exhibited more ¹³C-depleted eggs and less ¹³C-depleted fat bodies, suggesting increased lipid allocation to reproduction and reduced lipid storage.
